# Above- and below-ground links mediated by arboreal ants and host tree modify soil aggregation scaling, infiltration, and chemistry

**DOI:** 10.1101/2023.10.06.561289

**Authors:** Nicholas Medina, Lauren Schmitt, Ivette Perfecto, John Vandermeer

**Affiliations:** Ecology and Evolutionary Biology, University of Michigan, Ann Arbor, MI USA; School for Environment and Sustainability, University of Michigan, Ann Arbor, MI USA

**Keywords:** soil structure, invertebrate fauna, complex system, agroecology, above-belowground interactions

## Abstract

Soils are increasingly recognized as complex systems, emphasizing a need to study unique properties such as long-tailed scaling laws and the role of indirect interactions among arboreal above- and below-ground soil invertebrates. However, few studies consider the above-below-ground connections mediated by invertebrates’ activity and behavior compared to mediated by tree physiology. Given previous work showing that arboreal ants can compete and affect ground foragers as well as alter foraging behavior on different host trees, it is plausible that persistent above-ground ant nesting could extend to affect soil properties including structure and chemistry, mediated by ground ant exclusion. This study analyzes soil aggregation, water infiltration, and macro-chemical data associated with longer-term (5+ year) ant nesting in a rustic tropical agroforest. Results show that, 1) ant nesting maintained scaling law exponents or fractal dimensions of soil aggregate size distributions, and was significantly associated with relatively larger micro-aggregate diameters and log-normal variance in macro-aggregate size distributions, suggesting more consistent (less variable) underlying aggregation processes similar to host tree species effects; 2) areas in the vicinity of trees with no dominant ant nests had three-times faster water infiltration than in the vicinity of trees with dominant ant nests; and 3) a tendency toward changes in soil carbon and nitrogen stocks by one-quarter depending on host tree. These patterns are consistent with expected effects of ground ant suppression by an aggressive keystone arboreal ant, and are supported by previous studies reporting positive ground ant nest effects on soil chemistry and documenting ground ant foraging as a source of soil aggregate fragmentation. This study presents new ecological processes affecting ecosystem-scale functions, and suggests that future research on indirect interaction cascades would be beneficial to advance fundamental understanding of whole-ecosystem processes.

## Introduction

Soils are a basis for both ecosystem functioning (V. C. Yang, Papachristos, and Abrams 2019) and human civilizations (Montgomery 2007; Richter 2021). Accordingly, they are increasingly acknowledged as complex systems (Young and Crawford 2004) with many key interacting descriptive natural variables (Kuzyakov and Zamanian 2019), as well as less-studied social ones (Duarte-Guardia et al. 2020). However, both soil fauna and agroecological management (i.e. agroforestry polycultures vs. industrial monocropping) dimensions are continually under-studied (Bottinelli et al. 2015; Grandy et al. 2016) alongside soil microbial diversity and activity, prompting recent related syntheses (Lavelle et al. 2016; Barreto and Lindo 2022; Potapov 2022; Medina and Vandermeer 2023) and empirical efforts (Maaß, Caruso, and Rillig 2015). Specifically, ants are key soil engineers in the tropics (Griffiths et al. 2018), as earthworms are in temperate zones (Jouquet et al. 2006; Sánchez-de León et al. 2018; Angst et al. 2019), and in addition, themselves behave as a complex adaptive organismal system (Gordon 1996) whose collective soil engineering behavior is an extended phenotype (Jouquet et al. 2006) varying by taxon. Studying ant effects on soils also represents a unique coupling of naturally complex systems, especially in agroecologically managed agroecosystems where ant activity can be favored unintentionally (Vandermeer and Lin 2008). Furthermore, cascading effects to soils from aboveground ants in highly connected ecological networks (Vandermeer et al. 2019) can also represent a type of link between above and belowground processes, which remains a fundamental area of study in ecology (Bardgett and van der Putten 2014; Tao, Hunter, and de Roode 2017; Meier and Hunter 2018).

Analyzing soils using complex systems theory (Newman 2011) offers both perspective and interdisciplinary statistical tools including mathematical formulations with network structures (Chen et al. 2022), spatial explicitness (Vandermeer and Yitbarek 2012) for studying soil structure and heterogeneity (Rabot et al. 2018; Vogel et al. 2022), agent-based modeling (Waring et al. 2020), and novel parameters of non-normal skewed distributions, like scaling or power laws (Clauset, Shalizi, and Newman 2009), interpreted in part using fractal geometry (Boddy et al. 1999; P. Baveye, Parlange, and Stewart 2000; Menéndez et al. 2005). Previous reviews of how various metrics have been used in soils (Rieu and Sposito 1991; P. Baveye, Parlange, and Stewart 2000; Pachepsky and Hill 2017) explicitly point toward the need for further explanations of how relevant properties like scaling or power laws arise, as well as how they both reflect and are shaped by ongoing biological processes. The need for interdisciplinary interpretation may explain the ubiquity of more intuitive measures of soil aggregation like mean weighted diameter and water-stable aggregate proportion over others like fractal dimension (Caruso et al. 2011). However, fractal dimensions can offer useful, complementary, and more holistic representation of soil structured environments (Vogel et al. 2022), that are similar across soil phases and diverse components (Peng, Horn, and Hallett 2015), and also integrate combined effects of aggregate or pore size, density, and stability (Caruso et al. 2011) or related functions (Tisdall and Oades 1982), including implications for fine-scale biodiversity patterns (Alexandra N. Kravchenko et al. 2014; Rillig, Ingraffia, and Machado 2017; Bach et al. 2018), particulate and aggregate-associated organic matter storage (O’Brien and Jastrow 2013; Ananyeva et al. 2013; Alexandra N. Kravchenko et al. 2015; Alexandra N. Kravchenko and Guber 2017; Quigley and Kravchenko 2022), and local greenhouse gas emissions (Y. Li et al. 2019; J. Wang et al. 2019) and/or microbial co-metabolism (A. N. Kravchenko et al. 2019; Franklin et al. 2021) across hotspots of activity (Kuzyakov and Blagodatskaya 2015). Indeed, changes in scaling law exponents are generally considered indicators of more fundamental changes in underlying rules (i.e., ‘regime’ shifts) of a system, which for soils, can be shifts in relative aggregate formation versus fragmentation, such as those caused by more fungal agglomeration versus invertebrate burrowing, but also represents a generalized mechanism that can apply to cluster-forming systems more broadly such as networks. Beyond the physical limits of natural patterns (Halley et al. 2004), shared scaling patterns across biology and ecology have been generally interpreted as indicators of habitat complexity (Loke and Chisholm 2022), biodiversity maintenance (Ostling et al. 2004), entropy (Harte, Umemura, and Brush 2021; Klöffel et al. 2022), and comparable solutions to optimizing information flow or conserving energy during system changes or adaptation (West, Brown, and Enquist 1997). As a result, analyses of soils that draw from complex systems theory, such as fractal dimensions of soil aggregate size scaling or power law distributions, or similar long-tailed families including log-normal and exponential arising from partially hierarchical soil aggregation (Tisdall and Oades 1982; Melo, Figueiredo, and Filho 2021), continue to offer integrated information about how soils function and respond to key soil biota, like aggregate fragmentation by invertebrates during niche or nest construction (Maaß, Hückelheim, and Rillig 2019) and formation by associated microbial activity (Maaß, Caruso, and Rillig 2015).

Invertebrate soil ecosystem engineers are widely recognized for their fundamental roles in shaping ecosystems (Perfecto, Vandermeer, and Philpott 2014), soil habitats (Jouquet et al. 2006; Lavelle et al. 2016), biodiversity (Thakur et al. 2020; Lavelle et al. 2022), and related processes (McGlynn and Poirson 2012; Filser et al. 2016), although they are less studied compared to soil microbes (van der Heijden, Bardgett, and van Straalen 2008; Banerjee and van der Heijden 2022; Charlotte et al. 2022). Extended phenotype engineers (Jouquet et al. 2006), namely ants, have been shown to affect soil geomorphology (Whitford and Eldridge 2013) including: changes in texture and lower bulk density (Dostál et al. 2005; Cammeraat and Risch 2008); an increase in general microporosity (Tschinkel 2005), water infiltration (Cerdà, Jurgensen, and Bodi 2009) and water availability (X. R. Li et al. 2014); increased soil nutrient availability (Wagner, Brown, and Gordon 1997; Wagner, Jones, and Gordon 2004; Shukla et al. 2013; Kotova, Umarov, and Zakalyukina 2015; Sankovitz and Purcell 2022) but lower availability of metals (Gramigni et al. 2013); increased plant root and shoot growth (Farji-Brener and Werenkraut 2017); and recently, also increase microbial diversity (Delgado-baquerizo et al. 2019; Baker et al. 2020) and changed community structure, including even by non-fungal specialist ants (Lindström et al. 2019), in addition to fungal specialist (i.e. Attine) ants (Meyer 2013; Reis et al. 2015). Although, some studies present soils (compaction, granulometric composition, and pH) as being what affects ant biodiversity and community structure (Rocha-Ortega and García-Martínez 2018; Costa-Milanez et al. 2017), most other available studies support the reverse, ant modifications of soil properties (Shukla et al. 2013; Kotova, Umarov, and Zakalyukina 2015; Sankovitz and Purcell 2022), ultimately aligning with the context behind this study. More specifically, clearer evidence of ant effects on finer-scale soil structural properties, such as soil aggregate or pore size distributions, remains rare, but may be useful for interdisciplinary ecological models (Bennett et al. 2019). Ant effects on soil aggregate size distributions, which tend to have long tails and are thus tied to fractal geometry and self-organization (Young and Crawford 1991, 2004; Perfect, Rasiah, and Kay 1992), could be considered as resulting from skewed balances between positive formation and negative fragmentation processes, analogous to some early explanations of scaling laws in ecology (Macarthur and Wilson 1963; Triantis, Guilhaumon, and Whittaker 2012; Chase et al. 2019). This research gap presents an opportunity to assess how ants, and soil invertebrates more generally, affect soil system processes (Levin 1992).

Indirect or higher-order interactions are also increasingly recognized as important for ecosystem network dynamics (Werner and Peacor 2003; Perfecto, Vandermeer, and Philpott 2014) and biodiversity maintenance (Bairey, Kelsic, and Kishony 2016), and those tied specifically to keystone ant taxa have been shown to affect ecosystem stability (Vandermeer et al. 2021). Direct and indirect competition among ant taxa, occurring across above- and belowground plant, litter, and soil nesting sites, can generate persistent landscape-level spatial patterns in species occupancy (Vandermeer and Yitbarek 2012; Perfecto and Vandermeer 2013; Perfecto, Vandermeer, and Philpott 2014; Vandermeer and Perfecto 2020; Vandermeer et al. 2022). In some cases, this can lead to a halo effect, where some local competitive suppression and/or exclusion occurs within a limited range near a focal nest site, although coexistence may be maintained at a larger spatial scale (Ennis 2010; Ennis and Philpott 2017; Ennis, Perfecto, and Vandermeer 2023; Salinas, Vandermeer, and Perfecto 2019; Schmitt, Aponte-Rolon, and Perfecto 2020). Furthermore, similar to effects of leaf-cutting ants (Hudson et al. 2009; Sousa-souto et al. 2012), when ant-plant mutualisms are involved (Vandermeer and Perfecto 2019; Vandermeer et al. 2019), individual ant taxon foraging behaviors can be altered by the presence of extra-floral nectaries in tree canopies (Godschalx et al. 2015; Passos and Leal 2019; Nogueira et al. 2020), which can benefit the tree (Wagner and Nicklen 2010), but may replace other existing phloem-feeding insect-tending relationships nearby (Perfecto and Vandermeer 2006; Styrsky and Eubanks 2007). In cases where ants have very noticeable effects on their surrounding habitat (Morris, Vandermeer, and Perfecto 2015; Vannette, Bichier, and Philpott 2017; MacDougal 2019; Wildtruth and Perfecto 2023), even arboreal nesting activity has been shown to be associated with litter communities (Donoso et al. 2013) and decomposition (Schmitt and Perfecto 2020), soil nutrients (Clay et al. 2013; Godschalx et al. 2015; J. M. Lucas, Clay, and Kaspari 2018), soil microbiomes (J. Lucas et al. 2017), and host trees (Livingston, White, and Kratz 2008). Surprisingly, these effects show ecological variation by individual colony or diet, similar to co-evolutionary specialization like in Attine ants (Schultz and Brady 2008) and termites (Nobre et al. 2011). Ultimately, nesting and activity patterns of aboveground keystone ants have the potential to extend belowground to affect soil properties.

This study analyzes potential above-belowground throughline linkages between a tropical keystone arboreal-nesting ant *Azteca sericeasur* and adjacent agroforest soil properties. The motivating hypothesis is that persistent *A. sericeasur* nests affect soil properties by out-competing other nearby ground-dwelling ants (Ennis 2010; Salinas, Vandermeer, and Perfecto 2019; Vandermeer et al. 2019), that would otherwise have positive effects on nearby soils, such as those previously mentioned including increased burrowing and thus opening soil macropore space. Specifically, it was predicted that soils under persistent *Azteca* ant nests would show evidence of less beneficial ground ant activity, namely: smaller parameters of long-tailed soil aggregate size distributions including scaling laws, reflecting weaker self-similarity in the soil aggregation process and potentially less efficient soil structural dynamics; slower soil water infiltration, due to lower porosity; and fewer available nutrients, due to limited water infiltration; when compared to soils not under ant nests. Furthermore, these *Azteca* ant nest effects would be mediated by host tree species, and be stronger near host trees without extra-floral nectaries, where competitive suppression of ground ant activity would be stronger.

## Methods

### Study site and system

This study took place at Finca Irlanda *(15.1732729, -92.3365757)*, a 300 ha tropical, organic, biodynamic, shaded coffee agroforest farm in the southern highland Soconusco region of Chiapas, Mexico. The agroforest is 950-1150 m.a.s.l. and gets ∼4500 mm MAP, with one six-month rainy season. The region is widely known for relatively high coffee production, with nearby regional soils including Phaeozems, Andosols, Luvisols, and Leptosols (international WRB system). On site, local soils appear as epidystric Luvisols (typically USDA Alfisols) (Gardi et al. 2015), indicating clay illuviation and higher base saturation relative to other soils in the group. Additionally, topsoils can show subangular blocky aggregates, and subsoils columnar structure with occasional fine red or yellow mottling *(Fig. S1a)*.

A permanent 45-ha long-term (∼20 year) census plot (see *Philpott et al.* 2008) has over 90 shade tree species, mostly uniformly distributed, except along roads (Vandermeer and Lin 2008). Most canopy trees are *Inga*, notably *I. micheliana*, which are maintained by periodic pruning (Philpott and Bichier 2012) as a Mayan cultural legacy (Valencia et al. 2015) and also later promoted by local extension agencies (Peeters et al. 2003) in part for nitrogen (N) benefits (Romero-Alvarado et al. 2002) from N-fixing bacterial root symbionts (Grossman et al. 2005). *I. micheliana* is notable for hosting both extra-floral nectaries and helmet scale (*Octolecanium*), which can also affect symbiotic ant foraging patterns (Livingston, White, and Kratz 2008). In contrast, *Alchornea latifolia*, the second-most abundant tree *(unpublished data)*, is not a legume and does not have extra-floral nectaries.

*Azteca sericeasur* is a local keystone arboreal ant *(Vandermeer et al. 2010; Vandermeer et al. 2019;* Vandermeer 2021) that nests relatively indiscriminately in shade trees in this census plot (Kevin Li et al. 2016), mostly in trunks or occasionally constructed carton nests (Clay et al. 2013). *A. sericeasur* foraging can extend to nearby coffee bushes within 10 m (Ennis, Perfecto, and Vandermeer 2023), having cascading effects on coffee plant-associated arthropods, via predation, competitive exclusion, and aggression (Vandermeer, Perfecto, and Philpott 2010; Perfecto, Vandermeer, and Philpott 2014; Vannette, Bichier, and Philpott 2017); via mutualisms with extra-floral nectaries and Hemiptera, like *Coccus viridis* (Hsieh et al. 2012; Hsieh 2015) and *Octolecanium* (Salinas, Vandermeer, and Perfecto 2019; Livingston, White, and Kratz 2008); and even have cascading effects on litter-dwelling ant communities (Ennis 2010; Salinas, Vandermeer, and Perfecto 2019; Schmitt and Perfecto 2020).

### Data collection

#### Nest tree selection and variables recorded

Sampling sites were chosen based on census records of trees with resident active *A. sericeasur* nests for at least seven years. Paired control sites were trees less than or equal to 50 m from a chosen nest without a recorded *Azteca* nest for at least five years *(Fig. S1b)*. Five nest-control pair sites were *I. micheliana* trees, and another five were *A. latifolia*, totaling 20 trees.

Ground slope, leaf litter depth, tree diameter at breast height (DBH, ∼*2.7* m), number of coffee bushes closer than three meters from the target tree, distance to nearest trail, and *A. sericeasur* activity, as the number of ants passing a fixed point on the trunk in one minute, were recorded for each tree.

#### Soil sampling

Four soil samples were taken around each tree one meter from the trunk base along the cardinal directions. Each sample was taken down to 10 cm depth using a 5.5 cm diameter bulb transplanter and collected in small plastic bags in 2018, and in 2019 with a 5.5 cm diameter slide-hammer corer *(404.05, AMS Inc., American Falls, ID)* lined with inner plastic sleeves.Each tool sampled similar volumes. Soils were oven dried at ∼50 C (over three to four days) and final weights was used to estimate bulk density.

#### Aggregate isolation

Each dried sample was sieved through a six mm wire mesh, and individual aggregates were carefully arranged equidistantly on a light-colored flat surface, similar to other approaches (Oades and Waters 1991), and the arrangement was digitally photographed. Photos were visually color-corrected to detect separate soil aggregates and analyzed for pixel cluster area measurement using the color-threshold settings *(Fig. S1c)*, followed by the *‘particle size analysis’* function in the public Java-based software *ImageJ v1.52a* (NIH, <u>imagej.nih.gov</u>), with the resulting tables exported and composited by tree identity (i.e., individual).

Sites were re-sampled in 2019 to amass ∼500 g soil, which was sieved to 6 mm, and cores were composited by tree site and further sieved through a stack of metal wire meshes 4000, 2000, 500, 250, 125, and 63 µm wide, shaken for two minutes at ∼100 oscillations per minute over five cm radius by hand to standardize across samples and resemble other studies, and remaining size fractions weighed.

#### Infiltration and chemistry

To assess infiltration potential, subsamples of unsieved soil were placed into infiltrometers made from 2-inch-wide PVC pipes, wire mesh, and plastic collection cups *(Fig. S1d)*. One-quarter liter (250 mL) water was added to each column, and two measures were recorded: the time for 100 mL to pass through the column, as an initial rate, and how much total water collected after five minutes, converted to a final rate.

Soil aggregations greater than six mm (collected on the largest sieve) from a subset of sites were composited by tree ID, and to align with previous chemical analyses at soil aggregate-level (not shown), the 10 largest aggregates, plus 10 smaller aggregates weighing ∼20 g, were homogenized by mortar and pestle and sent for Olsen available phosphorus (P) analysis (Olsen et al. 1954; Horta and Torrent 2007) *(ECOSUR, San Cristóbal, Mexico)* in 2018, plus organic carbon (C) and total nitrogen (N) analysis using a LECO Trumac CN combustion analyzer *(LECO Corp, St Joseph, MI)* in 2019.

### Statistical analysis

Analyses were primarily done in R version 4.2.3 (2023-03-15) (R Core Team 2023) after ImageJ and described below. All dependent variable measurements were tested for both separate and interactive associations with ant nests and host shade tree species using a linear mixed effects model, to improve effect detection power given a paired-site design (X. Yang et al. 2014). Two fixed effects were ant nest presence and host shade tree species, and random effects were site pair number along with co-variates of ground slope angle magnitude, leaf litter depth, host tree diameter, number of nearby coffee bushes, and distance to nearest used walking path. Significance was set at *alpha*=0.05 and marginal significance at 0.05 < *alpha* < 0.1, and only models that met residual normality and equal variance assumptions using Shapiro-Wilk normality and Levene homoscedasticity tests were used.

Soil aggregate size distributions were analyzed for scaling law fits based on goodness-of-fit tests and bootstrapping methods (Clauset, Shalizi, and Newman 2009). Briefly, size frequencies were accumulated, scaling function x-min and exponent parameters were estimated for the long-tailed region, and bootstrapping was done to assess how likely the slope parameter was, given the data. The same was done assuming other similar long-tail distributions, namely exponential, log-normal, and Poisson families, and these were followed by pairwise comparisons testing if fitted scaling laws with given exponent and x-min parameter values described the data either better or equally well compared to similar alternative distribution families. Pre-analysis, large macro-aggregation (>6 mm) size values generated from image analyses were filtered for viable sizes based on the empirical mesh size used, and converted from area to diameter by calibrating the smallest viable aggregate detected with the sieve mesh size. Micro-aggregate data was initially converted from mass to frequency assuming spherical aggregate shapes, and that the mass proportion of each sieve fraction to about half the collected soil mass was similar to the mass fraction’s proportion of soil by volume, which generally aligns with other methods assuming constant aggregate density (Perfect, Rasiah, and Kay 1992). However, to minimize any potential compounding calculation effects on resulting distribution parameters, original soil aggregate mass fraction data were ultimately used for distribution parameter estimations to minimize data transformations (O’Hara and Kotze 2010).

Nutrient values were converted to stocks by multiplying raw percents by bulk density as typically done, and given a significant correlation between carbon and nitrogen values, these data were bound together into a MANOVA and tested for fixed effects of ant nest presence, host tree species, and paired site. Phosphorus and carbon-nitrogen ratios were tested separately with a similar linear mixed-effects model with paired site as a random effect. A general ordination was done using all dependent variables above as well as environmental co-variates except fitted Poisson parameters and measured ant activity using a PERMANOVA with ant nest presence, host tree species, and paired site as fixed effects. Analysis code was produced usingR packages *here* (Müller 2020), *poweRlaw 0.70.6* (Gillespie 2015), *rstatix* from *tidyverse* (Wickham et al. 2019), *lme4 0.70.6* (Bates et al. 2015) and *lmerTest 3.1.3* (Kuznetsova, Brockhoff, and Christensen 2017), *vegan* (Dixon 2003), *bookdown* (Xie 2023), *grateful* (Rodriguez-Sánchez and Jackson 2023), and stored at <u>github.com/nmedina17/azteca</u>.

## Results

### Aggregation

All soil aggregate size distributions followed a scaling law among both micro- and macro-aggregates, with exponents mainly ranging between 1.5 and 2 *(Fig. 1)*. Nearly all sample soil aggregate size distributions were better described by the scaling law function family than by the Poisson function family. While scaling laws did accurately describe the data, nearly all samples could also be described by log-normal and exponential functions, which share long-tail qualities with power functions but need more parameters defined *(Table S1)*.

**Figure 1:**
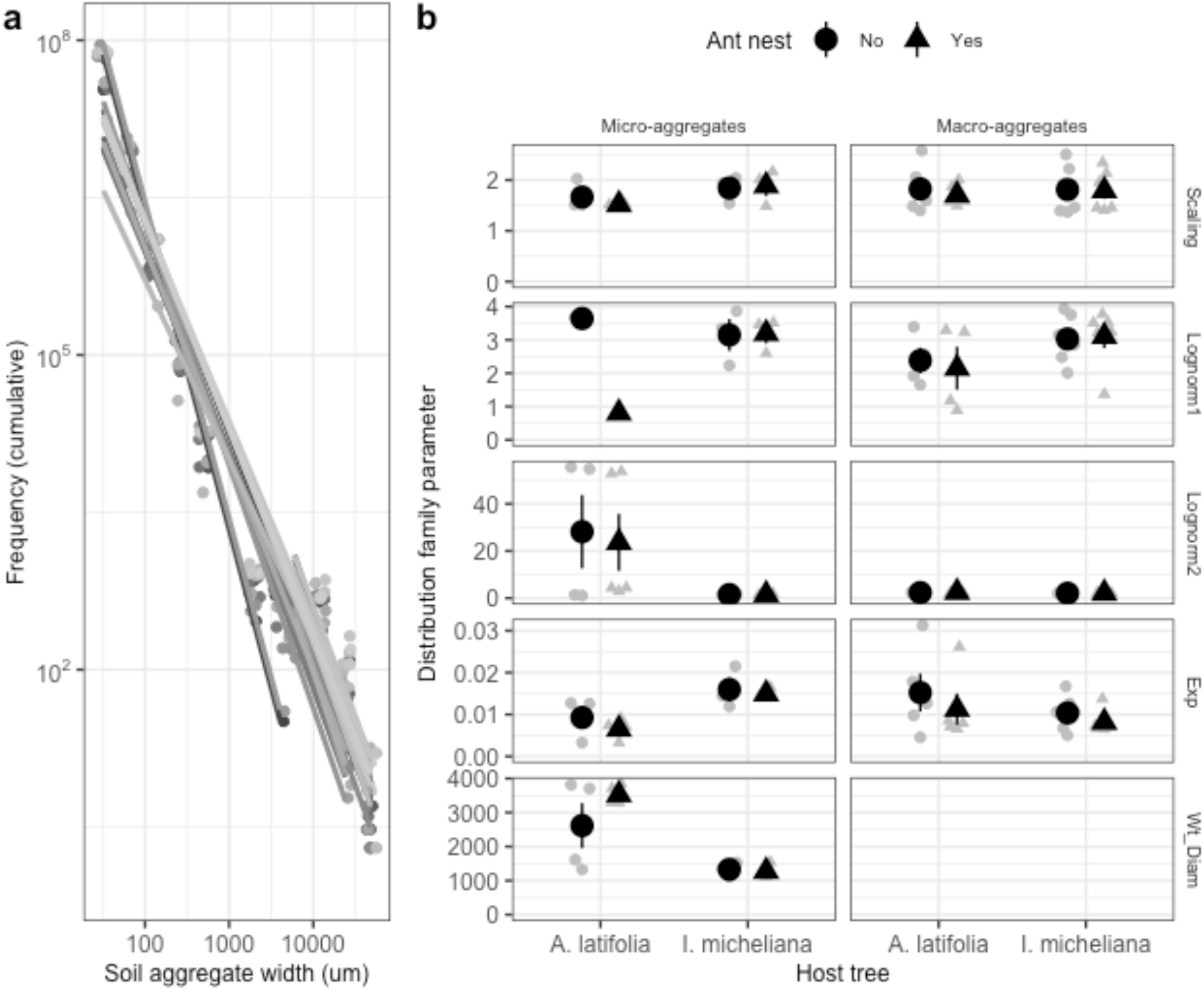
Soil aggregate size distribution (a) cumulative frequencies (log-transformed to show long tails) colored by local tree IDs where soils were sampled, and (b) long-tailed function family raw values (gray) and center means ±1 standard error (black) grouped by ant nest presence, categorized by host tree species, and facetted by both aggregate size class and distribution family.

Ant nests were not significantly associated with scaling law exponents (Fig 1b), but ant nests were significantly associated with 0.35 ± 0.13 units lower variance in macro-aggregate size, under a log-normal distribution model *(df=11.797, t=2.7, p=0.019*). Ant nests also marginally significantly increased mean weighted diameter by 903.82 ± 455.38 um *(df=6.2, t=<0.0001, p=0.092) (Table S2)*.

Host shade tree species was significantly associated with micro-aggregate size distribution parameters under all relevant models tested – scaling law, log-normal, and exponential function families. Specifically, micro-aggregate scaling law exponents were marginally significantly 0.33 ± 0.15 units lower under *Alchornea* than under *Inga (df=9.6, t=2.1, p=0.061)*, and associated exponential function base value parameters were 0.01 ± 0.0017 lower under *Alchornea* than under *Inga (df=5.9, t=5.2, p=0.0021)*. Both micro-and macro-aggregate central tendency parameters (but not variance parameters) under log-normal model representations were also lower under *Alchornea* than under *Inga*, by 1795.46 ± 107.74 for micro-aggregates *(df=0.91, t=17, p=0.049)* and 1.17 ± 0.48 for macro-aggregates *(df=6.8, t=2.4, p=0.046)*. Finally, mean weighted diameter of soil aggregates were 2248.97 ± 526.84 g larger under *Alchornea* than under *Inga (df=11, t=<0.0001, p=0.0014)*.

### Infiltration

Overall, soil water infiltration rates were significantly explained by both ant nesting and host shade tree species, both separately and interactively *(Fig. 2)*. Ant nesting alone significantly increased soil water infiltration by 5.54 ± 0.84 mL per sec initially after water entry and by ∼ 0.05 ± 0.01 mL per sec at near-complete infiltration (i.e., after most water had passed through the soil column).

**Figure 2:**
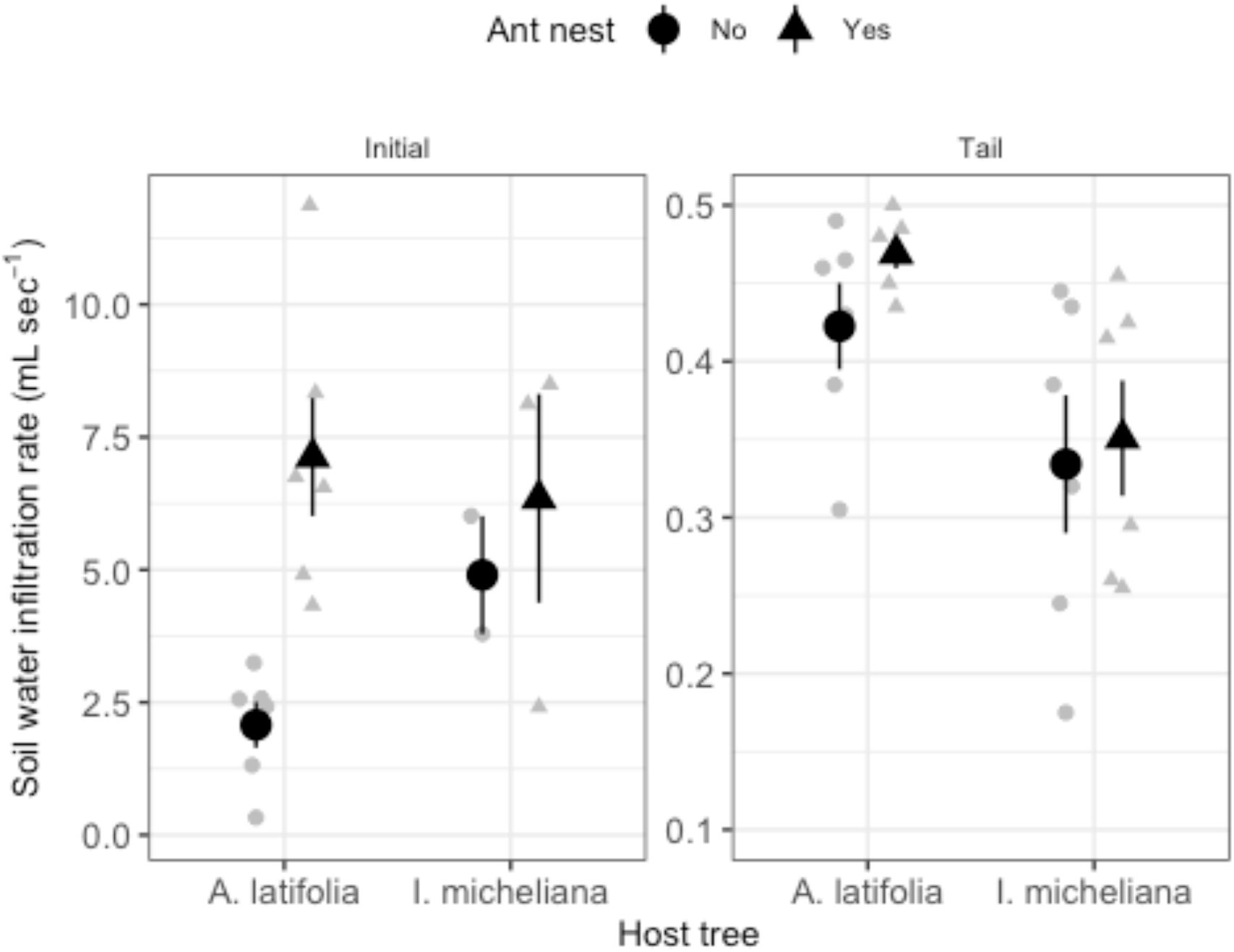
Soil water infiltration under *Alchornea* and *Inga* trees with (triangles) and without (circles) persistent (5+ year) arboreal *Azteca* ant nests, showing raw (grey) and center means (black) ± 1 standard error, during initial stages when water entered the soil column (left) and tail (final) stages of water percolation after most water exited the soil column (right).

Host shade tree species were significantly associated with water infiltration rates only at near-complete drainage, with *Alchornea* increasing rates by 0.11 ± 0.04 mL per sec *(df=13, t=<0.0001, p=0.028)*. Nest-tree interactions marginally significantly explained 4.74 ± 2.35 mL per sec of initial soil water infiltration rates *(df=12, t=2, p=0.066)*, which appeared as clear increase under ant nests on *Alchornea* but not under *Inga (Fig. 2)*.

### Chemistry

Overall, soil nutrients did not differ based on presence of ant nests but varied significantly among tree species and site *(Fig. 3)*. Soils showed lower nutrient stocks under *Alchornea* compared to *Inga* trees, specifically showing lower carbon significantly by nearly half *(df=1, F=28, p=0.0062)*, lower nitrogen significantly by also nearly half *(df=1, F=34, p=0.0043)*, and marginally lower phosphorus by 9 ± 4.2 *(df=8, t=2.2, p=0.062)*; there was no change in carbon-nitrogen ratio. There were also significant differences between host tree species by overall site pair (irrespective of ant nesting) in soil carbon *(df=4, F=9.1, p=0.028)* and nitrogen *(df=4, F=8.6, p=0.031)*.

**Figure 3:**
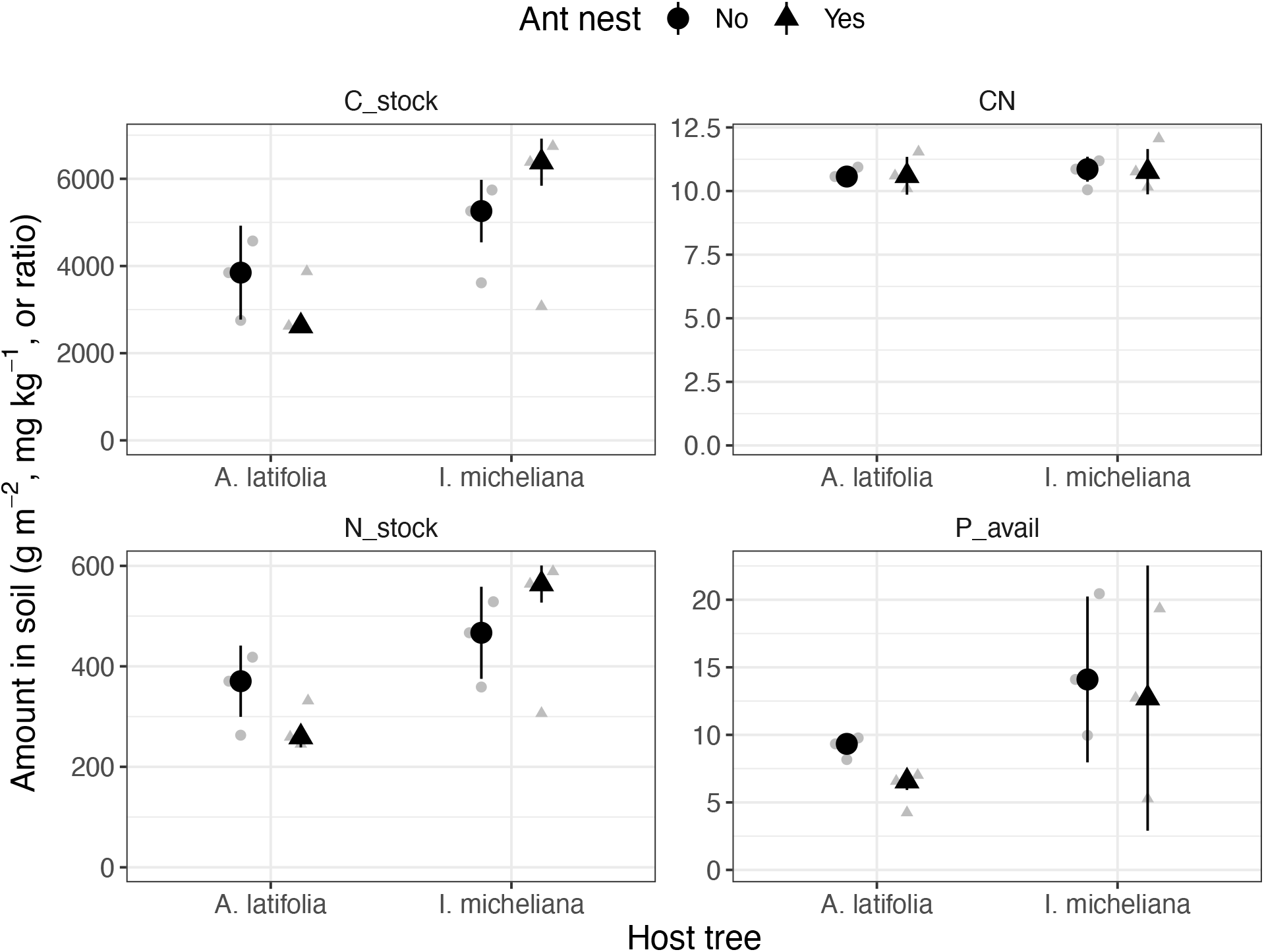
Soil carbon stock (top-left), nitrogen stock (bottom-left), available phosphorus (bottom-right), and carbon-to-nitrogen ratio (top-right) in soils under *Alchornea* and *Inga* trees with (triangles) and without (circles) persistent (5+ year) arboreal *Azteca* ant nests, showing raw (grey) and center means (black) ± 1 standard error.

### Co-variates

Including environmental co-variates, host tree species was significantly associated with ranked sample dissimilarities, explaining 13.8 % of variation in sample distances *(stress=0.089, df=1, F=3.5, p=0.044, R2=0.14) (Fig. 4)*.

**Figure 4:**
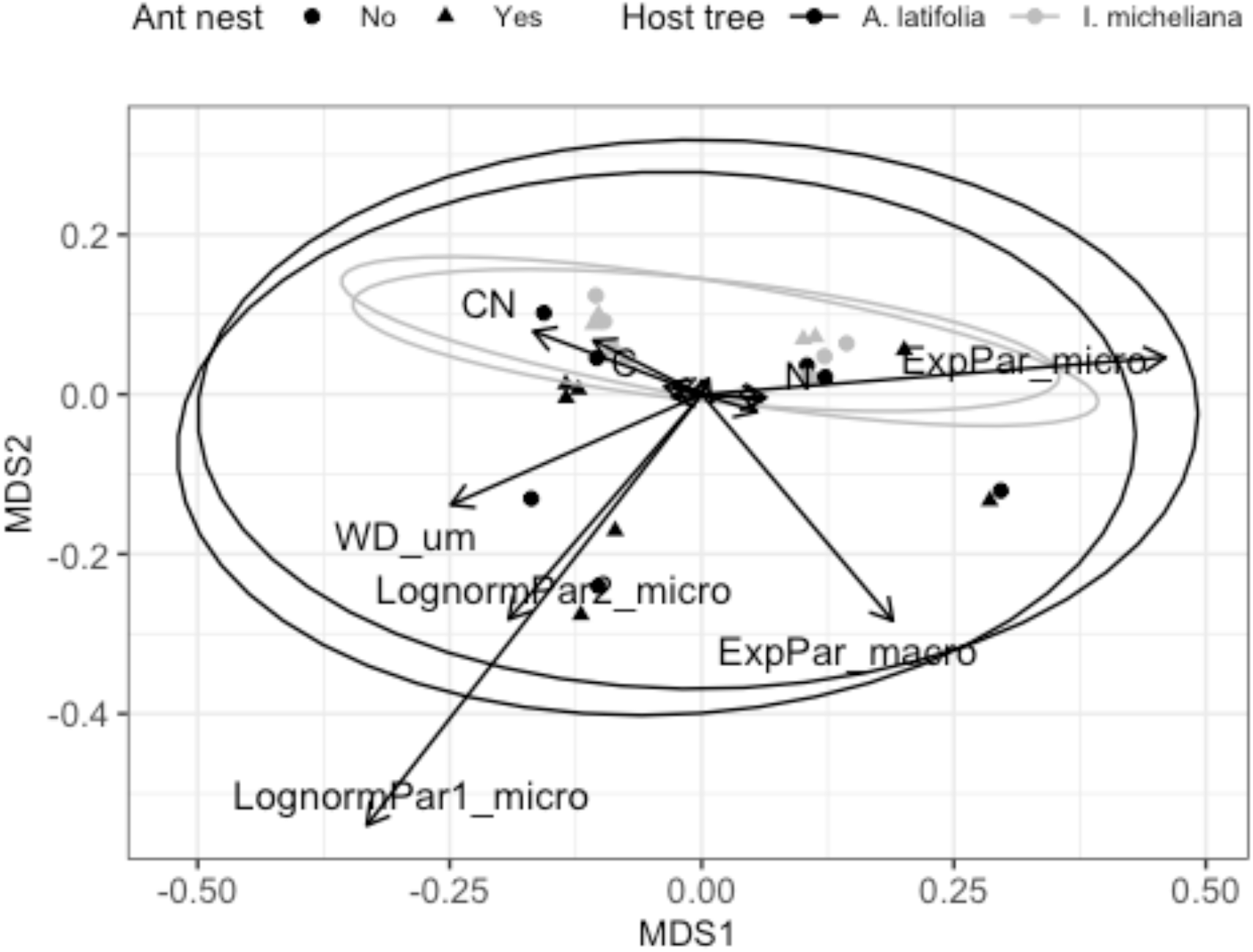
Plot of principle component analysis using all relevant numerical data grouped by points with (triangle) and without (circle) *Azteca* ant nests on non-leguminous *A. latifolia* (dark) and leguminous *I. micheliana* (light) shade trees in a coffee agro-forest.

## Discussion

Taken together, these results show that behavior-associated arboreal ant nesting patterns are associated with significant differences in soil structure, water infiltration, macro-chemistry, and environmental co-variates. Specifically, soil aggregate size distributions were described by distribution families with long tails, and their descriptive parameters were in various cases different across ant nest and host shade tree sites. *Azteca* ant nests alone marginally significantly increased mean weighted diameter and log-normal variance of soil aggregate sizes significantly. Alternatively, host shade tree *Alchornea* was associated with marginally lower micro-aggregate scaling law exponents, significantly lower exponential function bases and log-normal mean soil aggregate diameters, while increasing log-normal variance and mean weighted diameter – all of which suggests consistent soil aggregation producing relatively larger aggregates, generally improving soil structure. Soil water infiltration rates were also significantly higher under ant nests, marginally significantly interacting with host shade tree species. In line with faster infiltration, total soil nitrogen concentrations were also lower under ant nests, which may have been released in soil water. Overall, host tree species was a significant predictor of sampling sites across the focal tropical agro-forest. Collectively, results were somewhat inconsistent with initial hypotheses: *Azteca* ant nests were indeed partially associated with smaller soil aggregate distribution parameters, indicating relatively larger soil aggregates, which aligned with exclusion of ground ant burrowing and soil aggregate fragmenting activity; however, rather than infiltration being slower due to excluded ground ant burrowing activity, the opposing result of faster infiltration and lower soil nitrogen, potentially from leaching, aligned with the larger macro-aggregate results, which also suggested larger pores likely occurring between them. This suggests that arboreal ant nest effects on soils can appear direct, amid both potentially direct (Clay et al. 2013) and emerging indirect (Ennis 2010; Salinas, Vandermeer, and Perfecto 2019; Schmitt and Perfecto 2020) underlying processes.

Soil aggregation results here differed by response distribution parameter, suggesting that both arboreal ant nests and leguminous host shade trees with extra-floral nectaries can promote soil aggregation, but in different ways. Fundamentally, each metric of aggregate size distributions presented here reflects slightly different underlying processes. Scaling or power law distribution families by definition represent compounding multiplicative iterations among components governed by just one exponent parameter, also referred to as self-organization (Brown et al. 2002; Young and Crawford 2004), and can be generated by a preferential attachment process where ‘rich get richer’ (Newman 2001). In soils, this process could explain observed scaling laws if, as soil aggregates get larger, they also increase their chance of assimilating nearby smaller aggregates. Results here showed equivalent scaling exponents among macro-aggregates across both ant nest sites and host tree species, but not for micro-aggregates across host tree species, suggesting an interpretation that micro-aggregates were about one-third more likely to continue enlarging smaller, previously aggregated soil clusters into larger soil aggregates, thereby increasing self-similarity of all fine-scale soil aggregates, under leguminous nitrogen-fixing *Inga* trees, through two mm but before six mm. This faster scaling rate under a legume tree species could be explained by more fungal growth given higher nitrogen availability (Gou et al. 2023), since bacteria (Rashid et al. 2016) and fungi are well-studied soil aggregators (Ritz and Young 2004; Siddiky et al. 2012; Leifheit et al. 2014; Baumert et al. 2018; Lehmann and Rillig 2015; Lehmann et al. 2020), until soil aggregates get massive enough to be influenced more by plant roots than by thinner fungal hyphae (Rillig et al. 2015).

Additionally, it is increasingly recognized that scaling laws can also be generated by other processes (Simkin and Roychowdhury 2011), such as exponential functions composed together, which has been shown to also potentially represent exponentially distributed sampling times of components that themselves grow exponentially, as another related example mechanism underlying these long-tailed distributions (Reed and Hughes 2002). Exponential soil aggregation would reflect a relatively consistent soil particle accretion process, which is independent of aggregate size, as well as in this study ant nesting, but vary for micro-aggregates by host tree species. In line with a faster scaling law rate, there may also be some variation in size of smaller micro-aggregates used in the aggregation process, thereby equivalently explaining more fractional mass undergoing accretion under an exponential soil aggregation model.

Other long-tailed distributions such as log-normal distributions can also be generated by multiplicative processes, such as if the rates of multiplication or interaction strengths between components are themselves random and independently normally distributed (Mitzenmacher 2004). In practice, the log-normal peak or centrality parameter could also be thought of as translated into the start or minimum cutoff for scaling law shapes in empirical data (Clauset, Shalizi, and Newman 2009). Such an idea could help explain the significant host tree species effects on soil micro-aggregate log-normal centrality observed here, as could a possible underlying interaction with ant nest presence, but host tree effects on both log-normal centrality and variance in macro-aggregates suggests more change in process. Beneficial effects of nitrogen-fixing bacteria on fungal growth and soil binding may also be more variable in macro-aggregates, due to general increases in intra-aggregate heterogeneity, such as in hyper-local microbial community taxa and activity (Alexandra N. Kravchenko et al. 2014; P. C. Baveye et al. 2018; A. Kravchenko et al. 2019).

In soils, the aggregate hierarchy concept provides some theoretical support for such multiplicative exponential processes leading to long-tailed distribution families, given that different categories of processes from bacteria to plants roots are responsible for aggregating soil at different micro-spatial scales (Tisdall and Oades 1982; Melo, Figueiredo, and Filho 2021). In this case, each underlying processes such as clay flocculation, bacterial polysaccharides, and fungal enmeshment, would occur at a different rate. This is plausible, given that as soil aggregation proceeds hierarchically, the component sub-aggregates increase in mass and decrease in number, potentially slowing down the aggregation process, and is consistent with corresponding decreases in density and increases in porosity (A. N. Kravchenko et al. 2011; W. Wang et al. 2012) and heterogeneity (Barbosa and Gerke 2022) among larger macro-aggregates. Mean weighted diameter has also been discussed as an intuitive measure of soil aggregation (Caruso et al. 2011) and likely mathematically biased toward larger values (Pachepsky and Hill 2017). In cases where larger aggregates are more strongly affected, such as in this study for *Azteca* ant nests and shade tree species, this can be more likely to reflect significant group differences. In the field, relatively larger aggregates are consistent with the hypothesis of competitive suppression by aggressive ants such as *Azteca* (Ennis and Philpott 2017) of ground ant worker activity, whose mandibles fragment soil aggregates during tunneling and excavation for nest construction (Mikheyev and Tschinkel 2004; Tschinkel 2005; Cerquera and Tschinkel 2010).

Overall, considering scaling laws as the most parsimonious model with fewest parameters, in this study, soil aggregate size distribution exponents were around two, which agrees with image-based values of structural heterogeneity (Armstrong 1986; Perfect, Rasiah, and Kay 1992; Peyton et al. 1994; Kun Li et al. 2018), but less so soil water-based exponents reported from other studies (Tyler and Wheatcraft 1990, 1992; Kozak, Sokotowski, and Sokotowska 1996; Perfect et al. 1996; Perrier et al. 1996) and approaches (Young and Crawford 1991; Menéndez et al. 2005), barring methodological exponent calculation differences (Newman 2005). These exponents around two can be interpreted as when soil aggregates double in diameter, the chance of subsequently doubling again cuts nearly in half, and consistently across several orders of magnitude in soil aggregate size, as well as across varying ecological conditions. This process could either be extended to consider the whole soil mass and the pool of aggregates smaller or larger than a certain diameter, and/or zoomed in to consider that a single soil aggregate will typically be comprised of two slightly smaller aggregates, regardless of binding mechanism. Such a process in soils could be explained in a variety of ways, and warrants future research to improve fundamental understanding of universal ways that soils behave.

Faster soil water infiltration is also consistent with overall soil aggregation results of relatively larger soil aggregates, despite this study’s original hypotheses. Geometrically, larger soil aggregates make larger pores between them, allowing faster water flow and drainage (Franklin et al. 2021). Faster water infiltration could also occur due to optimal arrangement of soil aggregates, which also creates ideal mixes of macro- and micro-pores, and which optimally would increase connectivity and shorten longest path length among soil pore networks (Gastner and Newman 2006). Significantly faster soil water infiltration under *Alchornea* host trees, which do not have extra-floral nectary structures in the canopy, is consistent with suppression of ground ant activity. Similarly, unchanged water infiltration rates under *Inga* trees, which have extra-floral nectaries in the species used here, is consistent with un-altered ground ant activity, primarily as a source of soil aggregate fragmentation over formation. Taken together, this hypothesis was supported with a significant interaction between ant nest presence and host tree species, in the directions predicted by the original hypothesis. Slower infiltration under *Inga* could be explained by a more self-similar soil aggregation process, where more smaller aggregates are combined to produce aggregates of the same size, possible due to increased fungal enmeshment (Ritz and Young 2004), potentially increasing soil compaction.

Generally, soil nutrients, namely carbon and nitrogen, results tended to show some support for the hypothesis, and were consistent with results of other measured variables. Soils had less total carbon and nitrogen, and plant-available phosphorus under non-leguminous *Alchornea* host trees, where aggregates were observed to be simultaneously relatively larger and water infiltration was faster. This suggests that available elements may be more easily lost than stored in soil aggregates, despite the suggested importance of aggregates for organic matter stability (Johan Six et al. 2002; J. Six et al. 2002, 2004; Alexandra N. Kravchenko et al. 2015) and microbial diversity (Alexandra N. Kravchenko et al. 2014; Rillig, Muller, and Lehmann 2017). Alternatively for phosphorus, fungal scavenging and phosphorus provision may help stimulate nitrogen-fixation under *Inga* legumes (Püschel et al. 2017). While showing no detectable difference, ant nests appeared to exacerbate these effects, nearly showing even lower element concentrations under *Alchornea* where these *Azteca* ants forage more on the ground impacting the ground ant community, and slightly higher concentrations under *Inga* where they forage more in the canopy. These effects would agree with previous studies showing altered soil chemistry (Sankovitz and Purcell 2022) and heterogeneity (Wagner, Brown, and Gordon 1997) near ground ants, which *Azteca* excludes near non-leguminous host trees (Ennis 2010; Ennis and Philpott 2017; Salinas, Vandermeer, and Perfecto 2019). Ground ant effects on soils include higher leachable dissolved carbon and nitrogen and available base cations alongside lower organic carbon levels (Bierbaß, Gutknecht, and Michalzik 2015), possibly mediated by local soil microbiome changes (Dauber and Wolters 2000; Lindström et al. 2019). This evidence may suggest shade tree effects on soil nutrients may be present and possibly context-dependent (Romero-Alvarado et al. 2002) related to local invertebrate and/or microbial activity.

Overall, this study shows significant effects reaching from above- to belowground activity, mediated by persistent (5+ years) arboreal nesting and extended community effects of a keystone ant ecosystem engineer on different host tree species. This study highlights the importance of trait-mediated indirect interactions (Werner and Peacor 2003; Schmitz, Krivan, and Ovadia 2004) for soil and ecosystem functions, including above-ground linkages (Bardgett and Wardle 2003), and frames these effects using analytical tools from complex systems theory (Perfecto, Vandermeer, and Philpott 2014; Newman 2011) applied to soils (Young and Crawford 2004; Lavelle et al. 2016; Medina and Vandermeer 2023) affected by invertebrate ant activity (Hsieh et al. 2012). Future research may further incorporate indirect interactions, including functional (Martin and Isaac 2015) and behavioral traits, as relatively novel mechanisms unique to including soil fauna in model representations of ecosystem functioning (Grandy et al. 2016), with additional implications for land management, and potentially diversifying agro-pedogenic trajectories if such ecological management was to be applied at larger local or regional scales (Phillips 2017; Kuzyakov and Zamanian 2019).

## Funding

This study was funded by NSF DEB Award #1853261 as well as Tinker and Walls block field research grant awards from the University of Michigan Department of Latin American and Caribbean Studies, Ecology and Evolutionary Biology, and the School for Environment and Sustainability.

## Declaration of interests

No conflicts of interest declared.

## Author contributions

All authors helped conceive, design, and fund the study; IP and JV supervised; NM and LS collected and analyzed field and lab data; NM and LS wrote early article drafts; NM revised later drafts; all authors edited and approved final version.

## Acknowledgements

Thanks to peers in the Perfecto and Vandermeer labs for discussion of early drafts; and Gustavo, Flor and Miriam for field support.

## Data statement

Code stored at github.com/nmedina17/azteca.

## Supplementary information

## Methods

### Site

**Figure S1:**
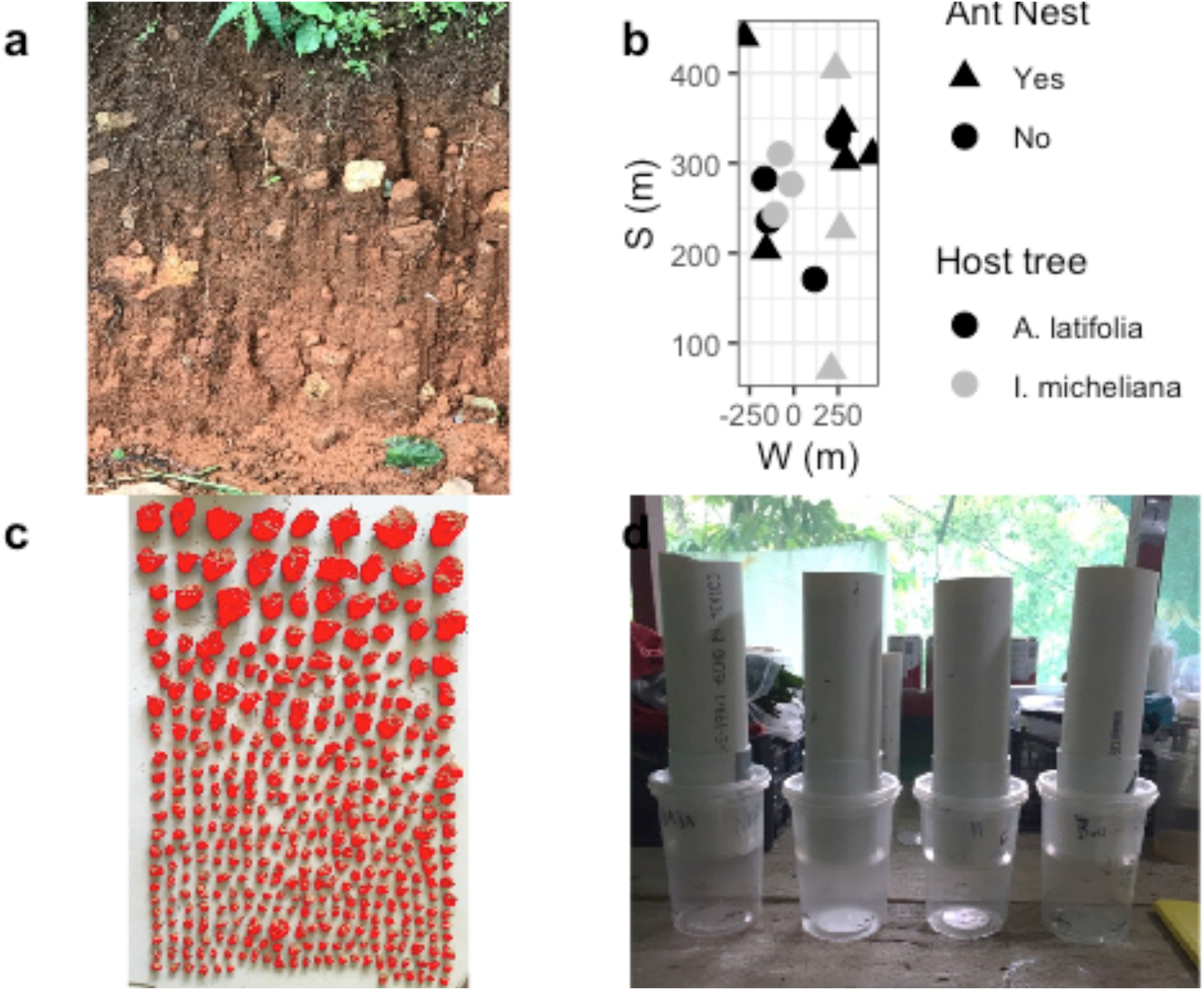
Field site and methods images, showing (a) road-side soil profile, (b) map of selected trees near sampling sites, (c) sample large macro-aggregate manual separation and cluster detection by ImageJ software (NIH), (d) constructed lab soil infiltrometers used in a remote diverse neo-tropical wet coffee agro-forest.

## Results

### Aggregate size distributions

**Table S1:**
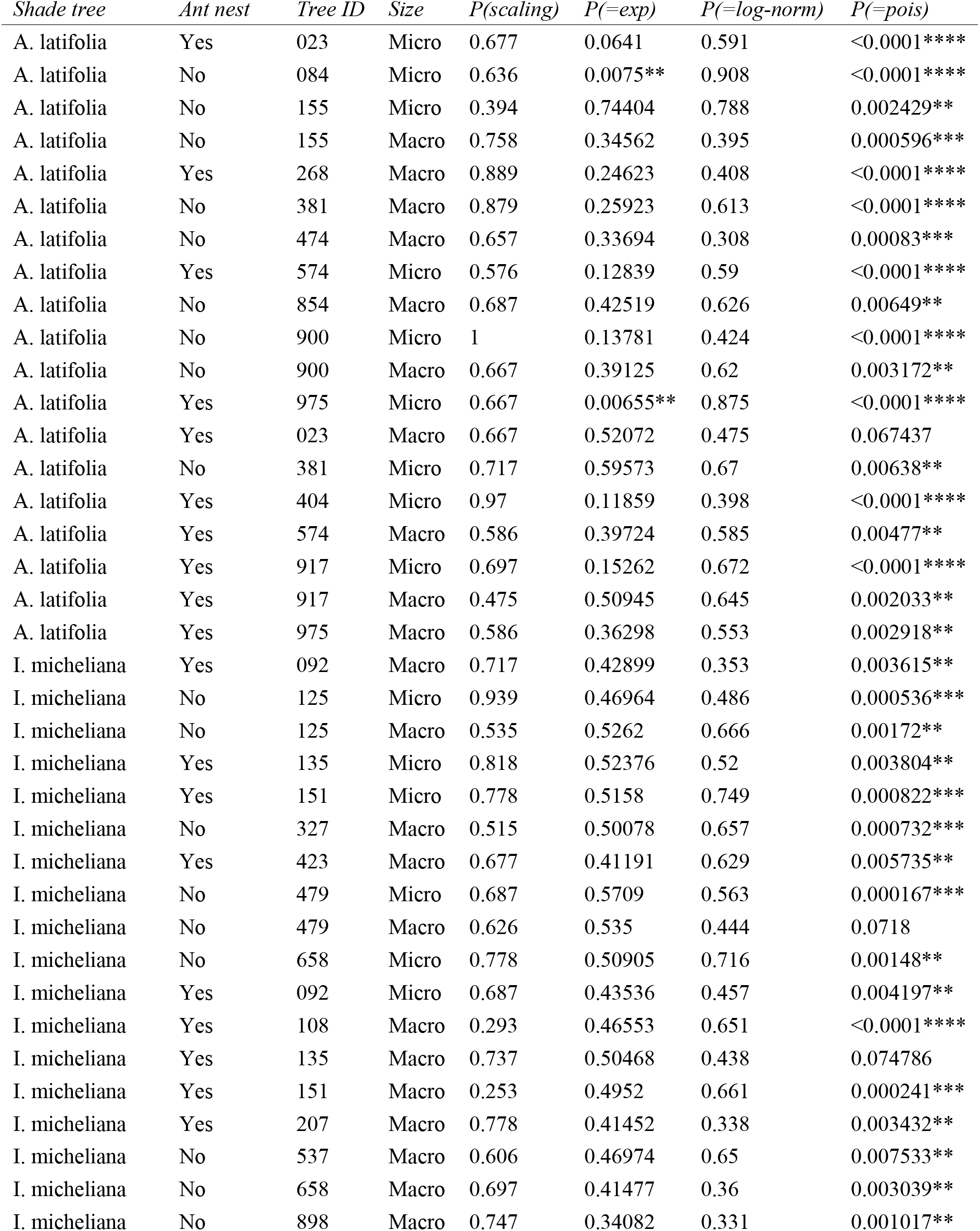
Scaling law fit and comparisons to similar distribution families for soil macro-aggregate samples. Null hypotheses H0 = sampled data follow a scaling law, and the scaling model is equally likely vs. comparison family.

### Statistical output

#### Aggregation and infiltration

**Table S2:**
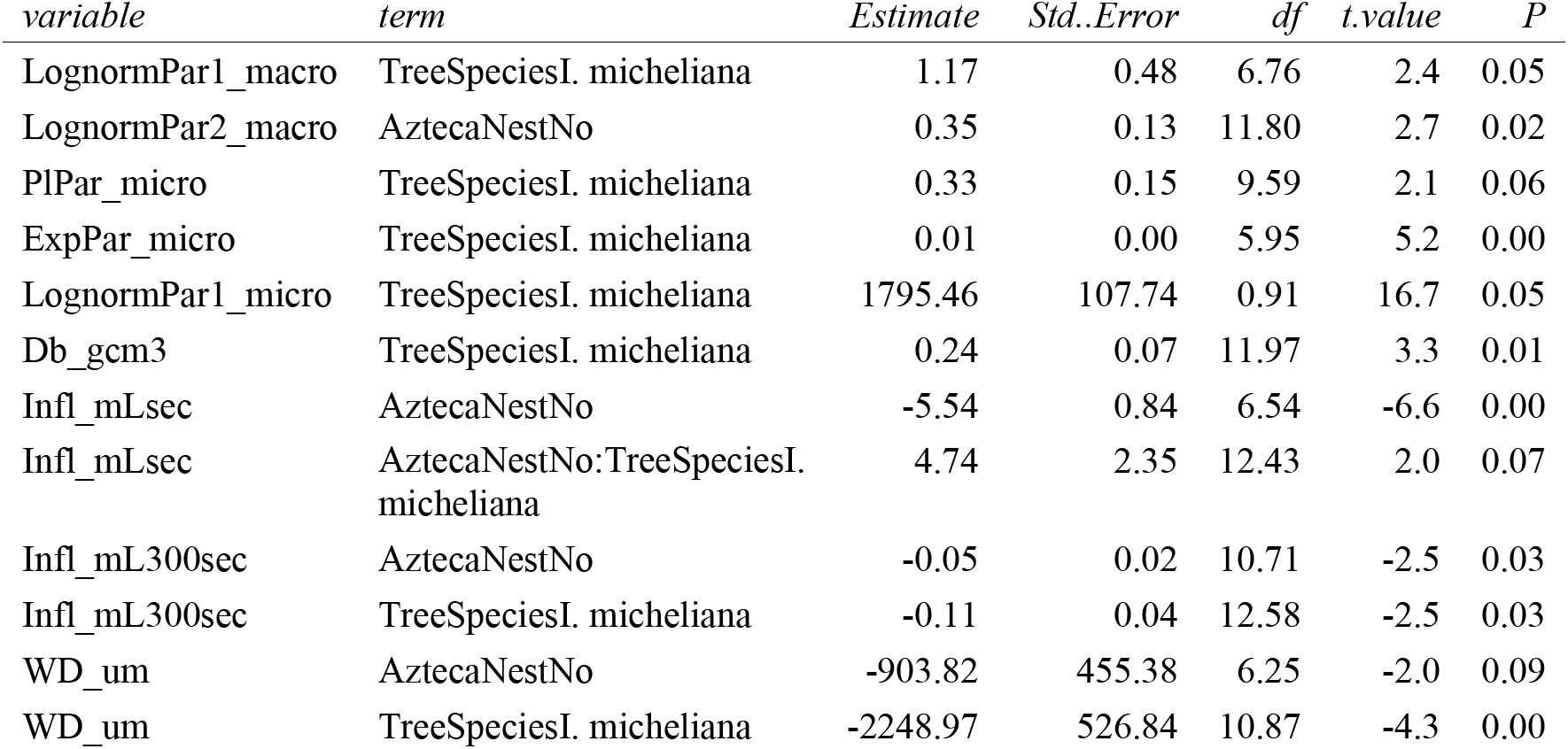
Statistical results for main soil aggregation and water infiltration data, using fixed nest and tree effects and random site pair and covariate effects.

#### Chemistry

Below: R code showing statistical model formula and result output for soil nutrient response variables.

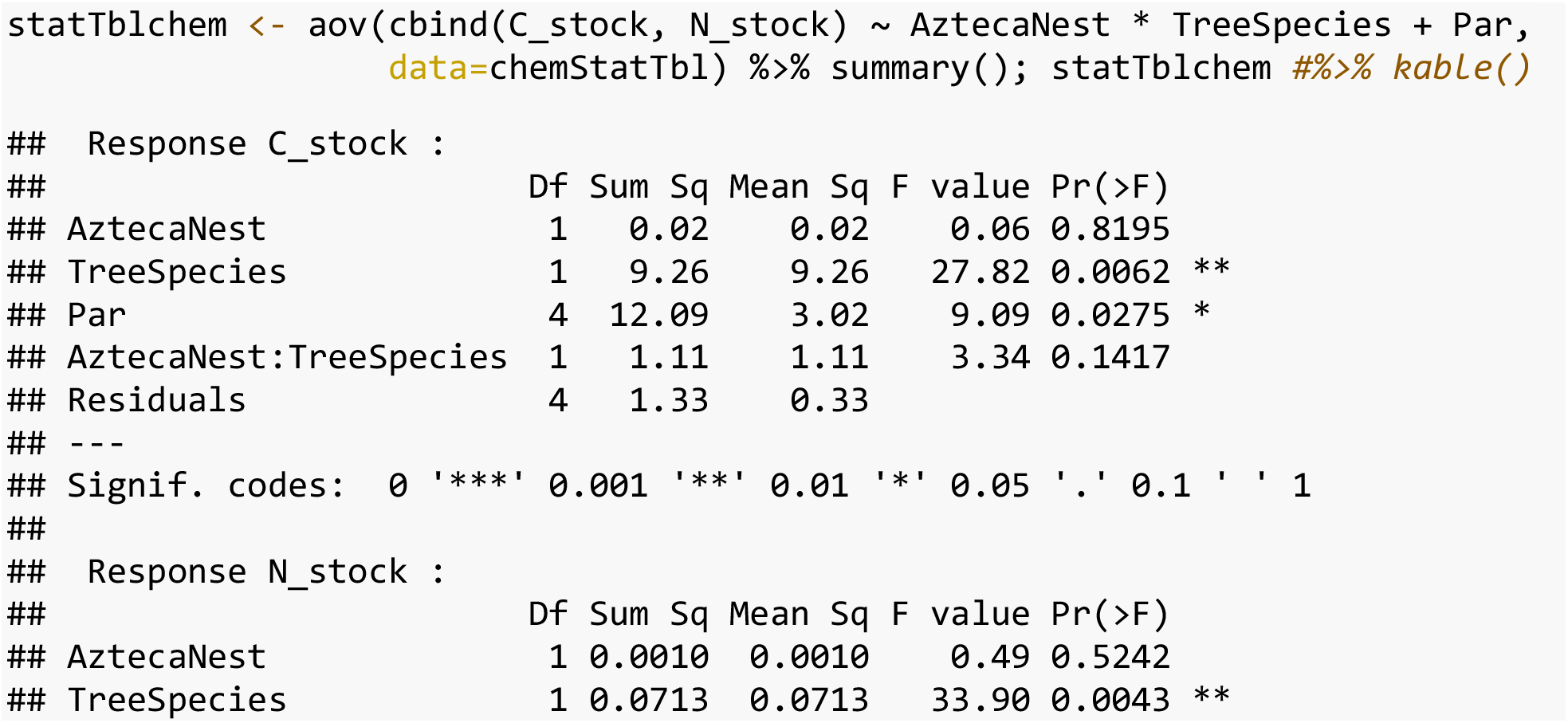
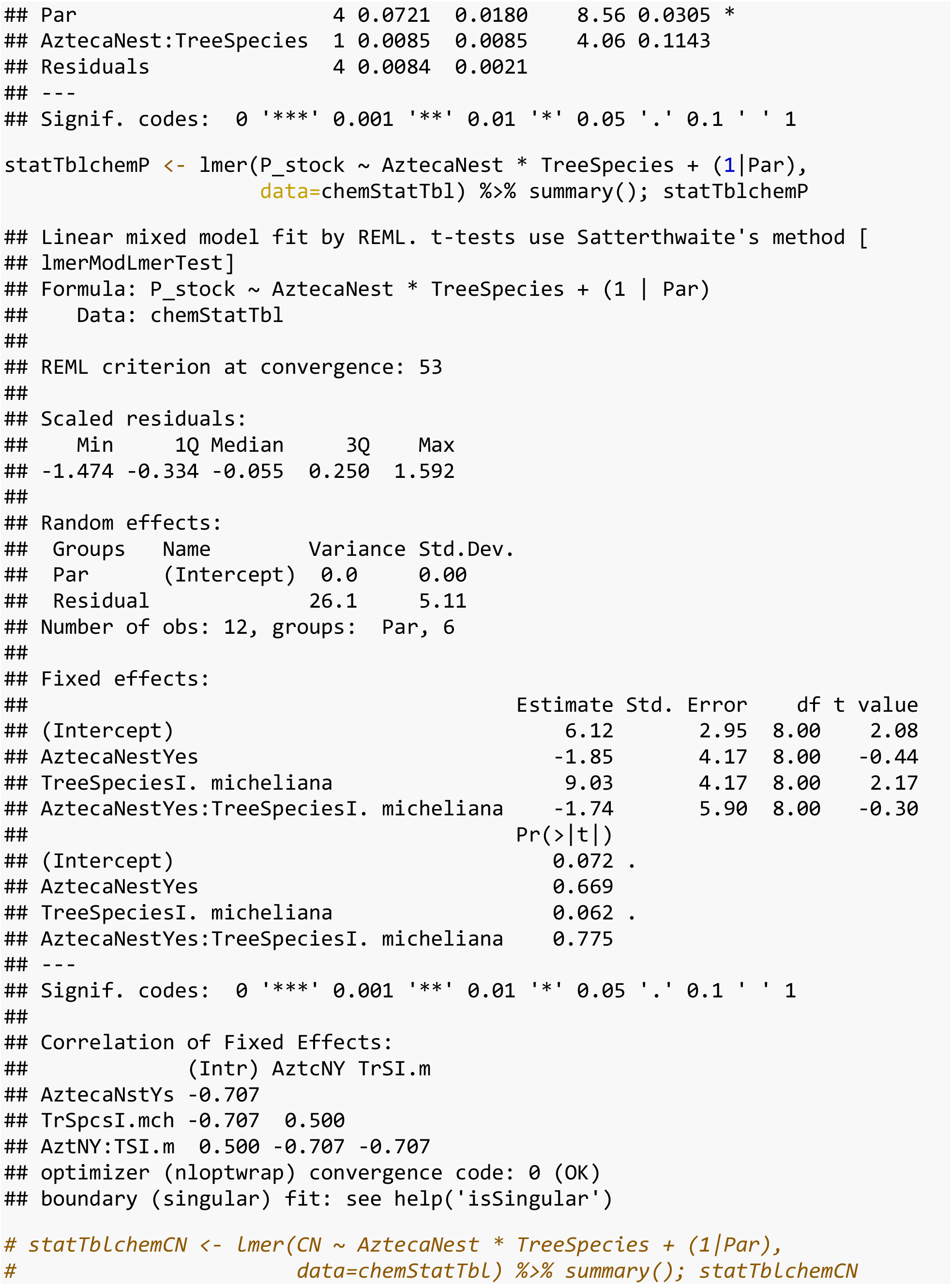

#### Co-variates

Below: R code showing statistical model formula and result output matrix ordination of environmental co-variates.

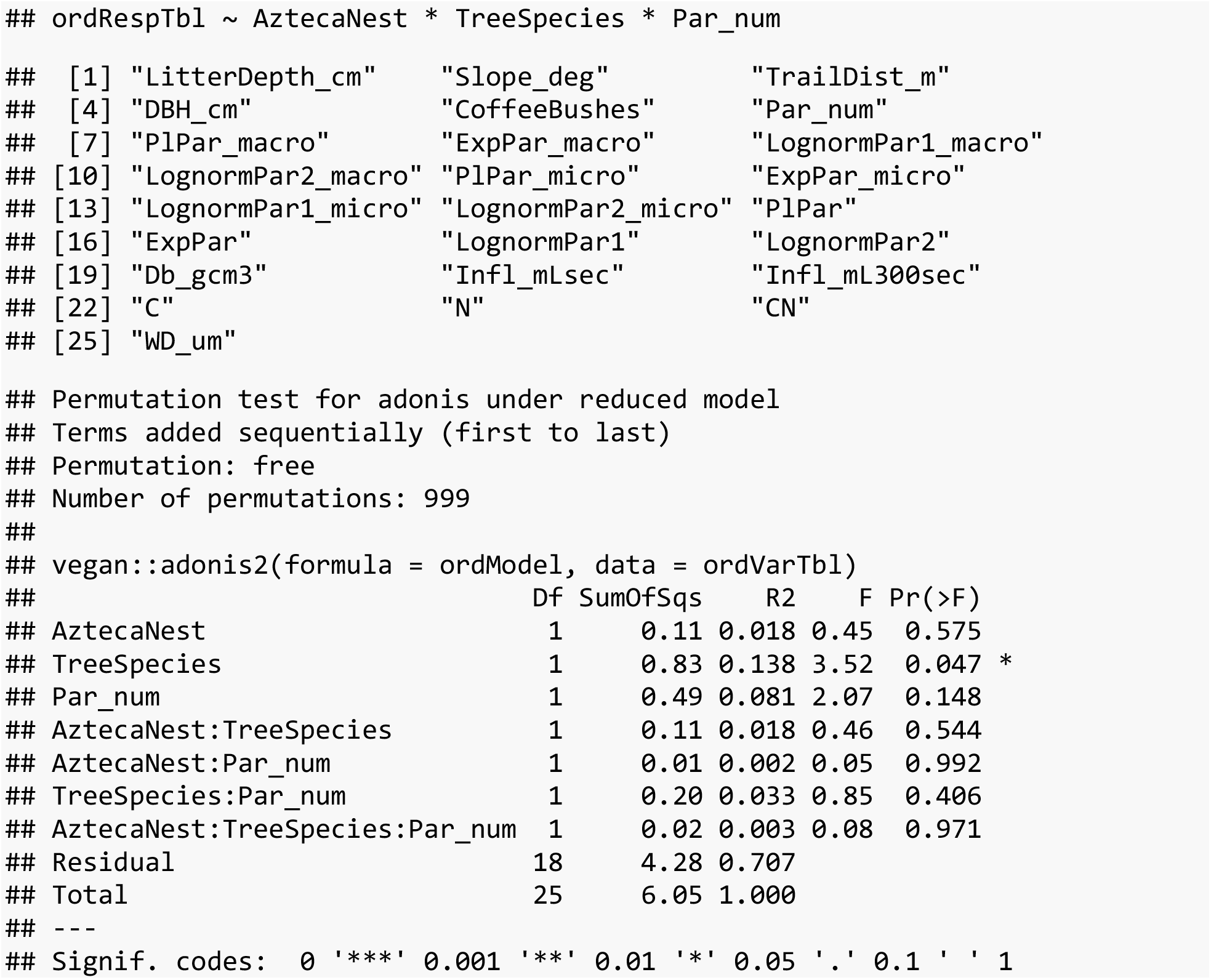

## Notes

### Competing Interest Statement

The authors have declared no competing interest.

